# Lactate transport at the uteroplacental unit- A theoretical study

**DOI:** 10.1101/2020.10.23.351841

**Authors:** Efrath Barta

**Affiliations:** Bar-Code Computers Ltd., Tirat-Carmel, Israel

## Abstract

Deciphering the lactate transport within the uteroplacental unit should be aided by a theoretical model in light of the insurmountable difficulties involved with in-vivo relevant measurements. Here we formulate a boundary value problem that predicts the direction and extent of lactate fluxes within the human placenta under various physiological conditions. It accounts for metabolic processes within the placenta and transporters’ activity at the two membranes that confine the terminal villi. Lactate concentration inside the terminal villi and its fluxes at the membranes are being computed. Under normal conditions lactate flux from fetal arterioles to the placenta surpasses the flux to the fetus via the umbilical vein. Within the placenta, it adds to the lactate that originates in the glycolysis, some of it degrades to pyruvate and surpluses are delivered to the maternal circulation. The apparent permeabilities of the placental membranes with respect to lactate as well as the specific characterizations of the placental lactate production, hitherto unknown, are being estimated. We determine the range of parameter values that induce sustainable, healthy fetal lactate levels and demonstrate the versatility of lactate exchange between the placenta and the fetus by computing the effect of extreme conditions (e.g., cesarean section, intrauterine growth restriction) on lactate fluxes.

## Introduction

Lactate is a vital metabolic substrate, consumed as an energy source in fetal heart, brain and skeletal muscles but also one that must be cleared when tissue hypoxia occurs. Basic facts regarding lactate transport in the human placenta remain elusive after decades of laboratory work, see the early perfusion experiments of Hauguel et al. [1], Medina et al. [2] and Schaefer et al. [3] that measured different directions of net lactate fluxes. Experimenting with the in-vivo fetus/placenta is of course inconceivable, animal placentas proved to be different from human ones [4, 5] and laboratory in vitro measurements are both very difficult to perform and have a limited utility. The placenta, the organ that allows nutrient shuffling between the maternal and fetal circulations, utilizes a rich variety of mechanisms in order to ensure a continuous supply and clearance of all vital substances. Diffusional exchange of metabolites occurs via the placental villi. Those are bathed in the intervillous space, IV which is filled with maternal arterial blood [6]. The placental villi are constructed as hierarchical trees with splitting branches, see Fig 1. The terminal villi, the smallest branches of the villous trees, are the main platform for nutrients exchange. They are being penetrated in their center by fetal capillaries so that the thin syncytiotrophoblast layer, ST that fills the terminal villi constitutes a barrier between the maternal blood in the IV and the penetrating fetal blood capillaries. Nutrients uptake and release occur at the membranes that confine those villi: 1. The microvillus membrane, MVM that wraps the villi. 2. The basal membrane, BM that wraps the fetal capillaries. Transfer of lactate at those membranes is determined not only by the concentration gradient of the monocarboxylate anion but also by the proton gradient due to activity of electroneutral, H^+^-coupled monocarboxylate transporters, MCT. Western blotting and immunolocalization techniques indicated at polarized MCT expression: the isoform MCT1 is more dominant at the BM while isoform MCT4 shows greater abundance on MVM. Those two members of the MCT family of symporters significantly differ in their kinetic characteristics [7, 8]. Within the villi, at the ST, glucose, pyruvate and lactate co-exist as glycolysis converts part of the glucose to lactate while via enzymatic reactions part of the lactate degrades to pyruvate.

**Fig 1.**
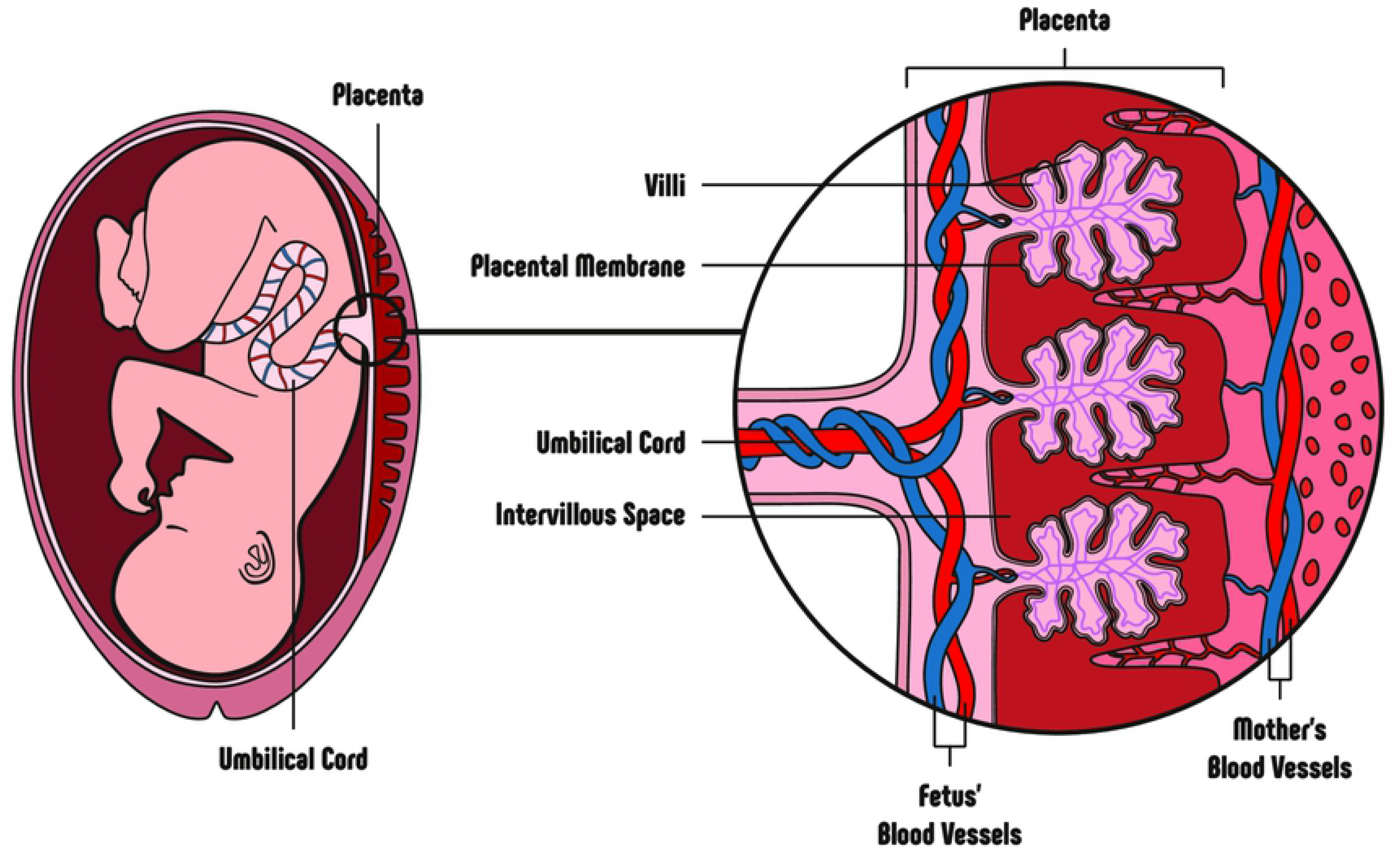
A human placenta anatomy diagram. The trees of villi are bathed in the maternal arterial blood that fills the IV and their “branches,” the terminal villi, are the main platform for nutrients exchange between fetal capillaries that reside within the villi and the maternal circulation. (This figure is right protected, *© **123RF.com***)

In contrast to other nutrients that are consumed by the fetus in a relatively steady manner, lactate transport might considerably fluctuate due to one or several of the following reasons: 1. Fetal lactate production or consumption rate is changed. 2. The acidity of the fetal and/or maternal circulation is changed. pH is determined by plasma concentrations of the strong cations (Na^+^, K^+^, Ca^+^, Mg^+^) and the strong anions (Cl^−^ and lactate) thus a significant change of any of them will affect the circulating plasma pH and consequentially, the diffusion rate at the confining membranes. Hence, the importance of a time-dependent simulation.

Here we formulate a theoretical model that accounts for the mechanisms that are involved in the bi-directional transport of lactate between the three components of the uteroplacental unit - mother, placenta and fetus, all are both lactate producers and consumers. It results with the unknown lactate concentration within the ST and the extent and direction of its fluxes that assure healthy, normal fetal lactate concentrations. Mathematically speaking, the model is a boundary value problem where relevant mechanisms that occur within the ST are described by a partial differential equation and the transporters-assisted diffusion via the membranes consist the boundary conditions. We found that placental lactate is mainly evacuated to the maternal circulation while its transport to/from the fetus is much more modest and, due to its high sensitivity, the net flow might switch directions in accord with the transient clinical situation. Lactate transport is shown to be determined mainly by MVM permeability and rate of glycolysis.

## Methods

Lactate fluxes within the uteroplacental unit are computed for given lactate concentrations and pH values in the maternal and fetal circulations.

### The governing boundary value problem

We formulate a model that refers to the mechanisms that govern synthesis, degradation and transport of lactate within the ST, the main barrier between maternal and fetal circulations, see Fig 2.

**Fig 2.**
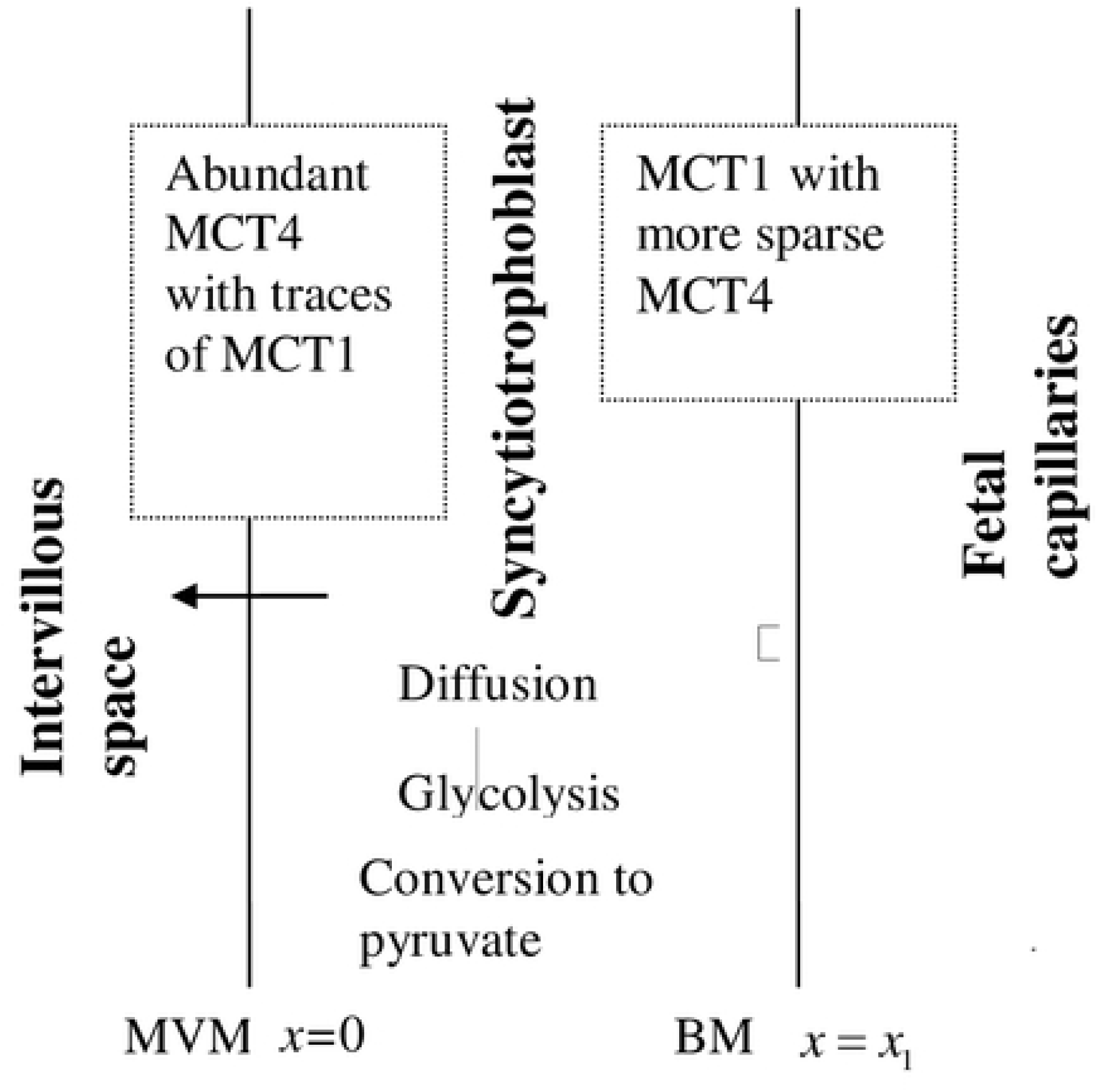
Placental lactate transport: 1D conceptual representation sketched not to scale. The region of our solution is confined by the two membranes that wrap (from outside and within) the terminal villous where lactate is produced by glycolysis, being degraded to pyruvate and moves along a concentration gradient. MCT1 and MCT4 transporters enable facilitated permeation at the membranes.

The high variability and complexity that characterize the placental villi tree as well as previous studies [9, 10] that computed a meaningful gradient of various substances’ concentrations just along the normal (to the membranes) direction, justify formulation of a 1D model. Within the ST, denote by *C_L_* the local, instantaneous concentration of lactate and by *H_p_* the uniform concentration (due to their high diffusion rate) of *H*^+^ ions. *C_L_* is the solution of Eq. 1 which accounts for the diffusion of the lactate, its production by glycolysis and conversion to pyruvate,both under aerobic conditions. Glycolysis is a multi-step metabolic pathway that yields two moles of lactate out of each mole of glucose and depends on the activity of quite a few enzymes. As the rate limiting glycolytic enzyme is Phosphofructokinase, known to be highly sensitive to the pH [11], we approximate the glycolysis kinetics by its activity. The reversible reaction between lactate and pyruvate is catalyzed by the cytosolic enzyme lactate dehydrogenase, LDH [12]. Therefore, we assume that both production and consumption of lactate are represented by enzymatic reactions and obey Michaelis-Menten kinetic:

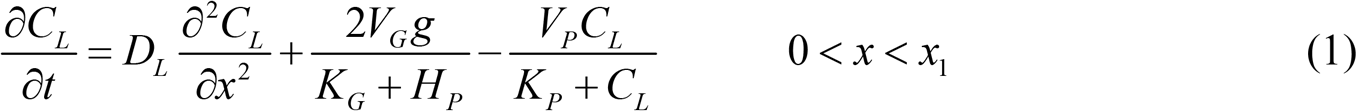

*x*_1_ is the thickness of the ST, *D_L_* is lactate diffusion coefficient in cytoplasm, *g* is the local glucose concentration, *V_G_*, *K_G_* and *V_P_*, *K_P_* are the kinetic characteristics of the reactions that govern lactate production and degradation respectively.

#### Boundary conditions at the MVM and BM

The density and affinity of the lactate-*H*^+^ symporters regulate lactate fluxes as passive diffusion via the two confining membranes was found to be negligible [13]. Turner [14] and later on Sanders et al. [15] have written lengthy expressions that explicitly describe the fluxes due to the action of symporters, entailing quite a few parameter values. In our case there are no laboratory established values of these parameters and we had to simplify the expressions in order to get estimation based on the available data, see S1 Appendix. At the MVM, the barrier between the placental ST and the maternal blood in the IV:

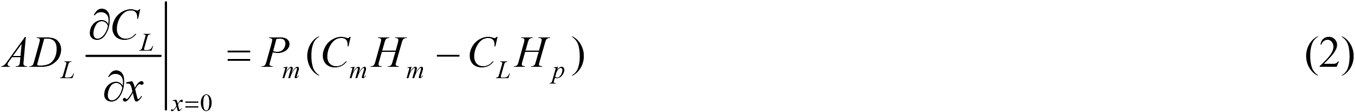

#### At the BM, the barrier between the placental ST and the fetal blood vessels (both arterioles and venules)

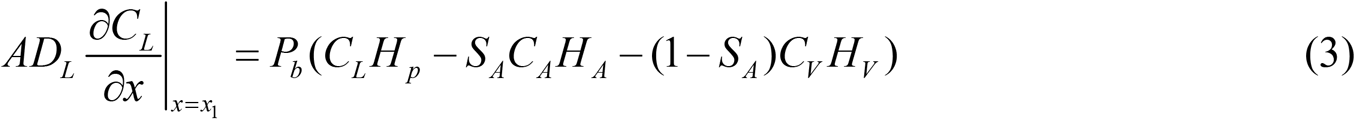

Here, *A* is the relevant membrane’ surface area, *S_A_* is the percentage of the arterial out of the total capillaries and sinusoids exchange area, *P_m_*, *P_b_* denote the permeability constants of the MVM, BM respectively. *C_i_*, *H*_*i*_ *i= m, A, V* are lactate and *H*^+^ concentrations respectively in maternal arteries, fetal arterioles and fetal venules. Note that we substitute in Eq. 2 arterial maternal pH and lactate level although lactate is delivered to the maternal circulation via veins that drain blood from the IV. This is because the blood in which the villi are bathed in the IV is basically an arterial blood, see Fig 1. Any initial condition for Eq. 1 will do as long as it is compatible with the boundary conditions.

### Calibration of the model (parameter values)

Parameter values are listed in Table 1. Here, we extend on their choice.

**Table 1.**
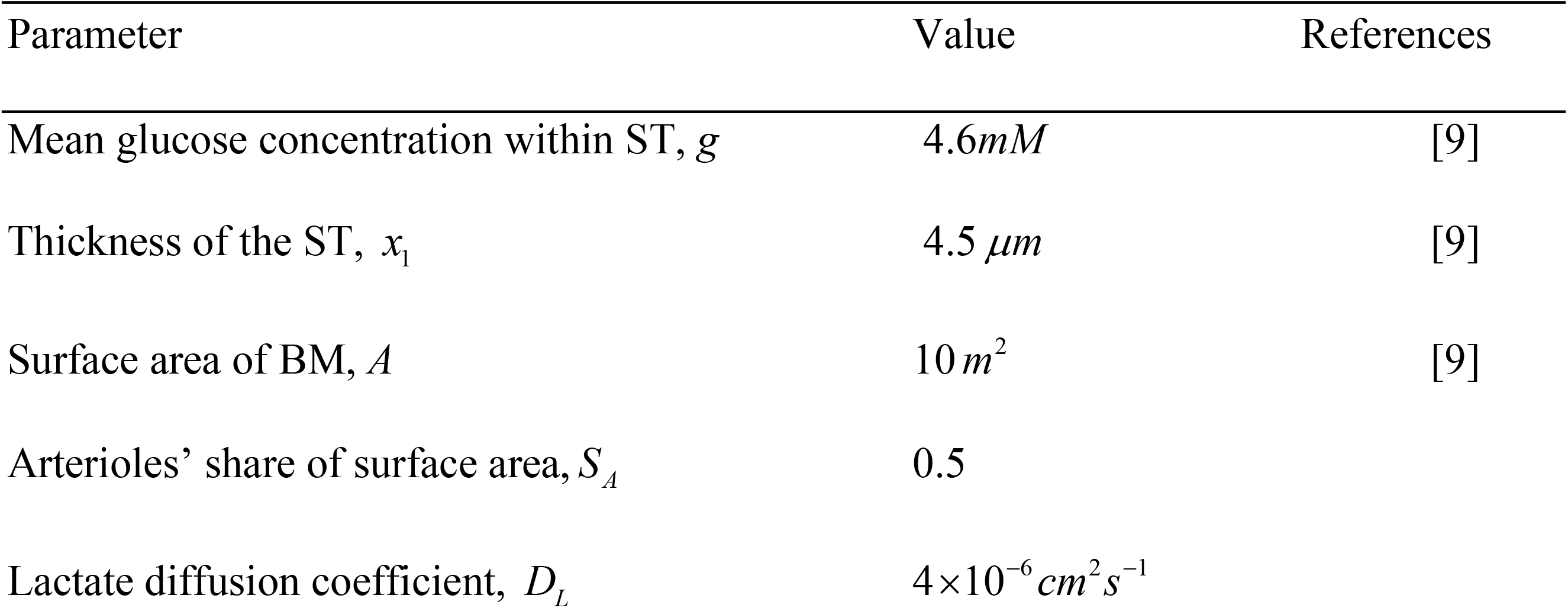
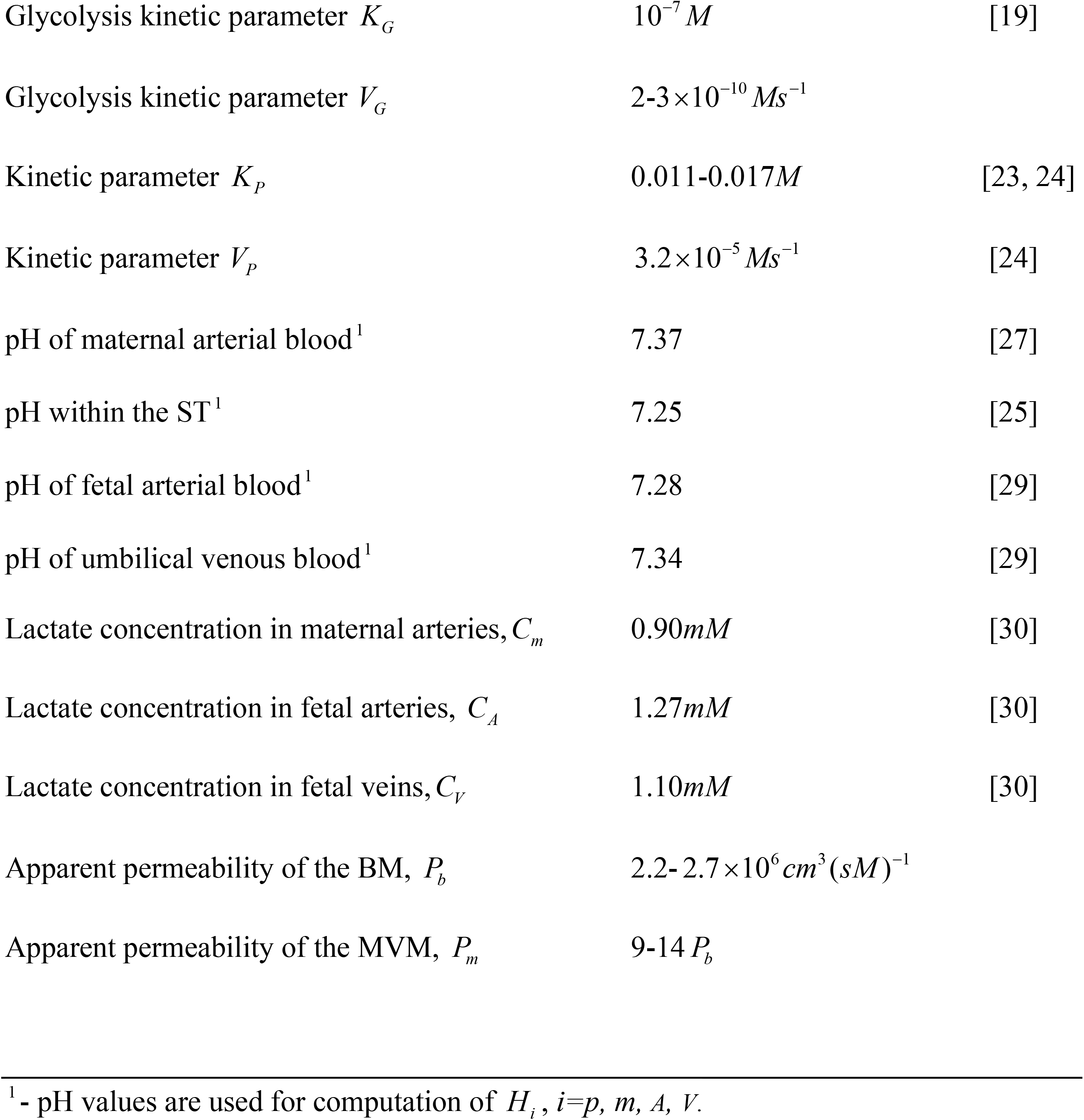
Literature-based and inferred values for our default set of parameter values

*g*: Glucose concentration varies just a little within the ST [9], thus for all practical purposes it has a constant value-about 4.6*mM*.

*x*_1_: The ST is a highly anisotropic layer with a thickness that varies from 1.5 *μm* to over 5 *μm*. We apply an averaged thickness [17].

*A*: Total surface area of the peripheral villi amounts to 12 *m*^2^ and surface area of the fetal capillaries and sinusoids is 0.7-0.9 of this value [9]. Here we use *A*=10 *m*^2^.

*S_A_*: In contrast to the situation in intermediate villi where arterioles are wider and occupy a more central position than venules [6], in terminal villi there is a huge variation between the location and size of the two types of the capillaries and sinusoids within the same placenta and between different normal placentas. Therefore, we use 0.5 as a mean value for *S_A_*.

*D_L_*: Diffusion rate of lactate in the cytoplasm is unknown. However, lactate molecular weight is half of the glucose weight and as diffusion rate is inversely proportional to the cubic root of the molecular weight, *D*_L_ amounts to 4.4×10^−6^ *cm*^2^*s*^−1^ based on the diffusion coefficient of glucose in the ST [9]. Another estimation of *D_L_* comes from measurements of lactate diffusion in the brain [18] that yielded *D*_L_ = 0.75-1.1×10^−5^ *cm*^2^*s*^−1^. Diffusion rate is inversely proportional to the viscosity of the ambient fluid and ST is 2-3 times more viscous than the fluids in the cerebral nervous system therefore, we deduce a value of about 4×10^−6^ *cm*^2^*s*^−1^.

*K_G_*: This constant is characteristic to the chemical reaction per se thus its value, 10^−7^*mol* / *l* is inferred from measurements in other organs [19].

*V_G_*: Bittner et al [20] found an order of magnitude difference between glycolytic rates in various types of cells thus *V_G_* is probably specific to the ST and cannot be inferred from measurements in other environments. Michelsen et al. [21] found that placental glucose consumption has a very high variation and on the average it amounts to 32 *μmol/min*. 40-70% of it is converted to lactate [1, 22] where each mole of glucose yields 2 moles of lactate. Glycolysis occurs within the ST - its volume approximately equals the product of *A* and *x*_1_. All these data result with a glycolytic rate (the 2^nd^ term in the right hand side of Eq. 1) of (10 –18)10^−6^ *M* / *s* that, after substitution of *g*, *K*_*G*_ *H_p_*, yields *V_G_*.

*K_P_*: Again, it is mainly a characteristic of the reaction thus we infer its value: 0.011-0.017*M* from measurements in other cells’ types [23, 24].

*V_P_*: Measurements in placentas are unavailable. We utilize a value cited in [24]. Note that it implies that under normal conditions about 15% of the lactate is converted to pyruvate. *H_i_*, *i=p, m, A, V*: pH measured in cultured human ST cells equals 7.25 [25]. Pregnant maternal arterial blood pH is 7.35-7.45 [26, 27]. Umbilical vein and artery pH equal 7.25-7.45 and 7.20-7.38 respectively [28, 29].

*C_i_*, *i=m, A, V*: We use the values measured by Marconi et al. [30]: arterial lactate concentration in pregnant mothers equals 0.90*mM* while the concentrations in fetal circulation are higher - about 1.10*mM* and 1.27*mM* in umbilical vein and artery respectively (concentrations are accurate within ± 0.2*mM*).

*P_b_*: Median cardiac output equals 425*ml/min* per kilogram of fetus [31]. About 40% of it is distributed to the placenta (and returns via the umbilical vein) [32]. Thus, 0.5-0.6 liter of blood is exchanged with the placenta each minute and in steady state Eq. 3 might be replaced by:

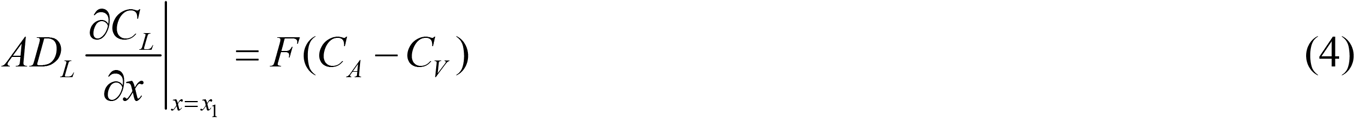

where *F*=550/60 *s*^−1^*ml*.

*P_b_* is obtained by equating the right hand side of Eq. 3 while using an estimated/guessed *C_L_* value and that of Eq.4 after substituting a 0.17*mM* concentrations’ difference [30]. We utilize this *P_b_* value to compute *C_L_* as the solution of the boundary value problem, Eqs. 1–3. In case that a non-negligible difference between the computed and guessed *C_L_* values arises, parameter values are changed iteratively till the guessed and the resultant *C_L_* values coincide, see S2 Appendix for more details.

*P_m_*: Western blots staining indicate that while MCT4 is abundant at the MVM with just traces of MCT1, MCT1 is dominant at the BM with non-negligible density of MCT4 there [16,8]. The exceptional density of MCT4 at the MVM compensates for its lower (compared to MCT1) affinity to lactate and explains the higher permeability of the MVM. However, western blots staining does not reveal the membrane permeability as it results with relative densities of transporters and not with absolute values. Moreover, the affinities of the membranes were measured per gram of membranes’ proteins where the mass of the MVM and BM were not estimated. Henceforth we implicitly infer the specific permeability constant from the available relevant data: Settle et al [8] measured a factor of 2.75 between the number of lactate moles per mg. of membrane’s proteins that cross perfused MVM and BM during 20 seconds. Taking into account the different proteins’ content in the 2 membranes [7, 33] the above 2.75 factor converts to 2.5 for fluxes per mg. of membrane. No data regarding membrane weights are available thus we have to fathom the ratio between those two weights based on these facts: MVM surface area is 6 times that of the BM [9] however its base is thicker (i.e., probably heavier) with protruding microvilli that are extremely delicate (therefore, probably much lighter). We assume that *P_m_* / *P_b_* =9-14. This is in accord with very early perfusion experiments that found lactate fluxes via the MVM to be about 9 times those that cross the BM [28].

### Numerical and analytical solutions

Toms731 solver of the JSim library applied to the boundary value problem Eqs. 1-3 yields time-dependent profiles of lactate concentration within the ST. Alternatively, one may analytically derive an approximation for the solution in steady state; after zeroing the left hand side of Eq. 1 and linearizing the differential equation by omitting *C_L_* at the denominator of the last term of the right hand side of the equation one gets:

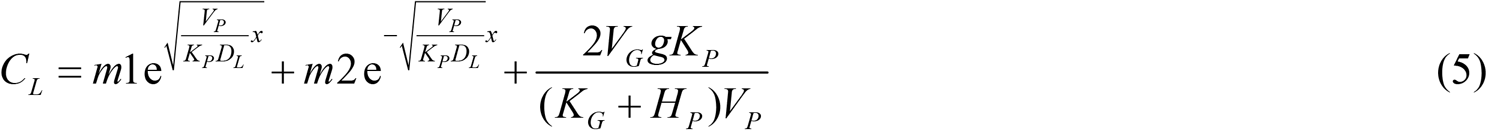

*m*1, *m*2 are found as the solution of the algebraic system of equations obtained by substitution of Eq. 5 in Eqs. 2,3:

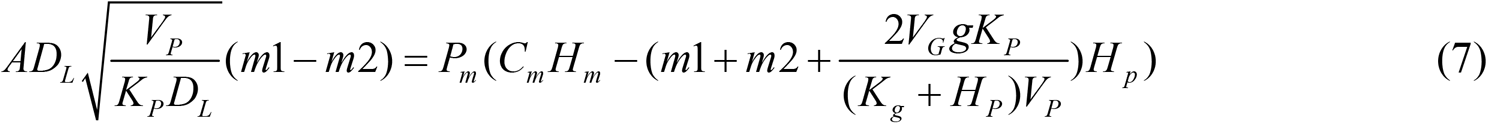

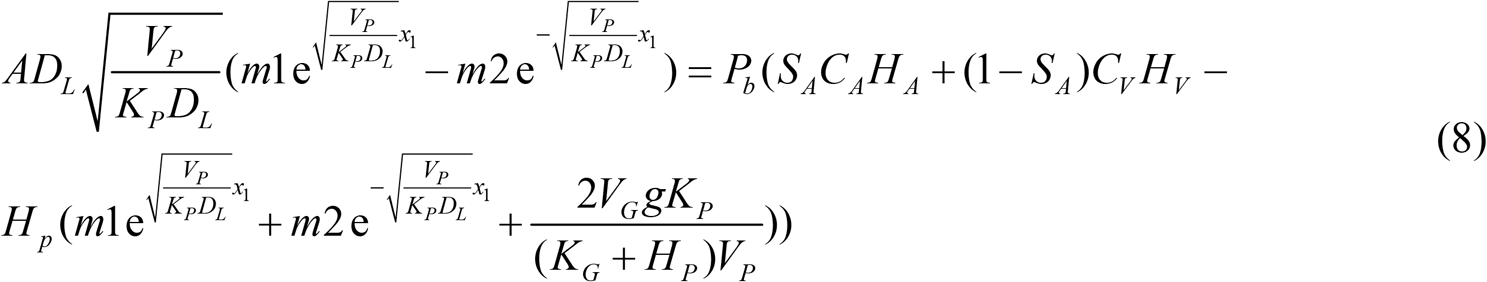

Steady state solutions obtained by the analytical and numerical approaches were found to differ by less than 0.5%.

## Results

Fig 3 demonstrates a typical situation where lactate flows from the fetus to the placenta and then to the maternal circulation. Placental lactate concentration (about 1*mM*) is in between maternal level and the higher fetal level. Significant changes of concentration occur at the confining membranes but it hardly varies within the ST. Nevertheless it would be a mistake to settle for solving a non-spatial problem where lactate concentration in the ST is assumed a constant value; the sensitivity analysis reveals the importance of the diffusion within the ST, see below, and in addition, ignoring local variations will obscure the difference between the regime of flow in the vicinities of the two membranes - the steeper slope near the MVM is an indication of the high rate of flow to the maternal circulation compared to the less significant fluxes that cross the BM.

**Fig 3.**
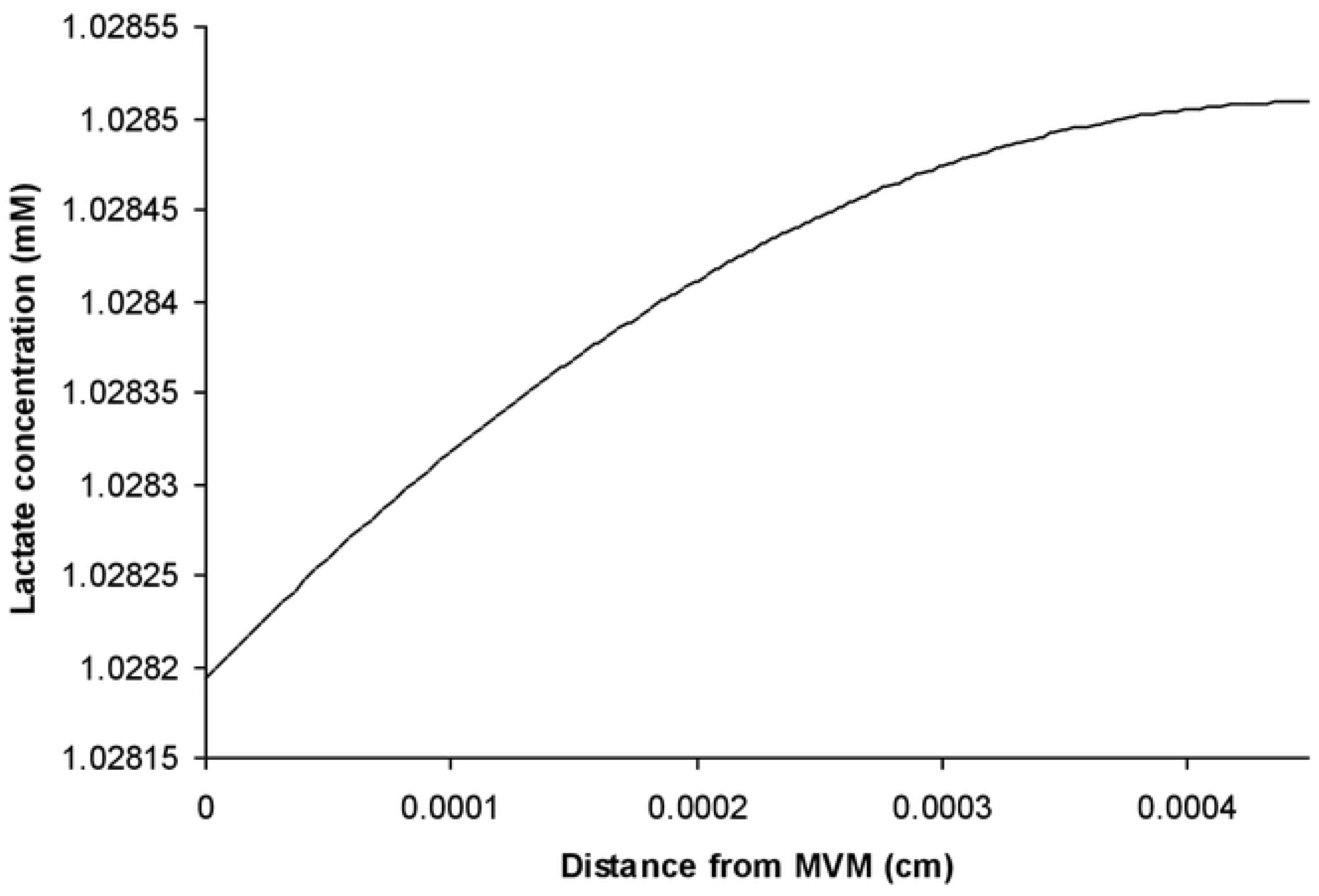
Lactate concentration within the ST as a function of distance from MVM. *K_P_* =0.013*M*, *V_G_* =2.5 ×10^−10^*M* / *s*, *P_b_* = 2.4 ×10^6^ *cm*^3^ / *sM*, *P_m_* =12 *P_b_*, other parameter values are specified in Table 1.

The main outcomes of this model are:

### A. Estimation of lactate fluxes at the two membranes under various clinical conditions

Some examples are specified in Table 2 – the first line represents our reference set of parameter values that corresponds to a steady state situation at the end of the pregnancy period when placental lactate is evacuated mainly to the maternal circulation and lactate outflow from the fetus is a bit higher than its inflow. This finding supports Marconi et al. [30] claim that the fetal hepatic uptake of lactate is its major route of disposal. Lines 2-4 of Table 2 demonstrate the effects of gradually changing the values of one or more parameters till a new steady state is established: A given increment of both arterial and venous fetal lactate levels is linked to 83% and 125% of that increment of placental and IV lactate levels respectively (line 2) mainly due to an enhanced lactate clearance at the BM. Thus the placenta in a way serves as a buffer that shields the fetus from a lactate overdose due to high maternal lactate level. Comparing lines 1 and 2 indicates that the same umbilical venoarterial lactate concentration difference might be associated with significantly different maternal lactate levels, in accord with past findings [34].

**Table 2.**
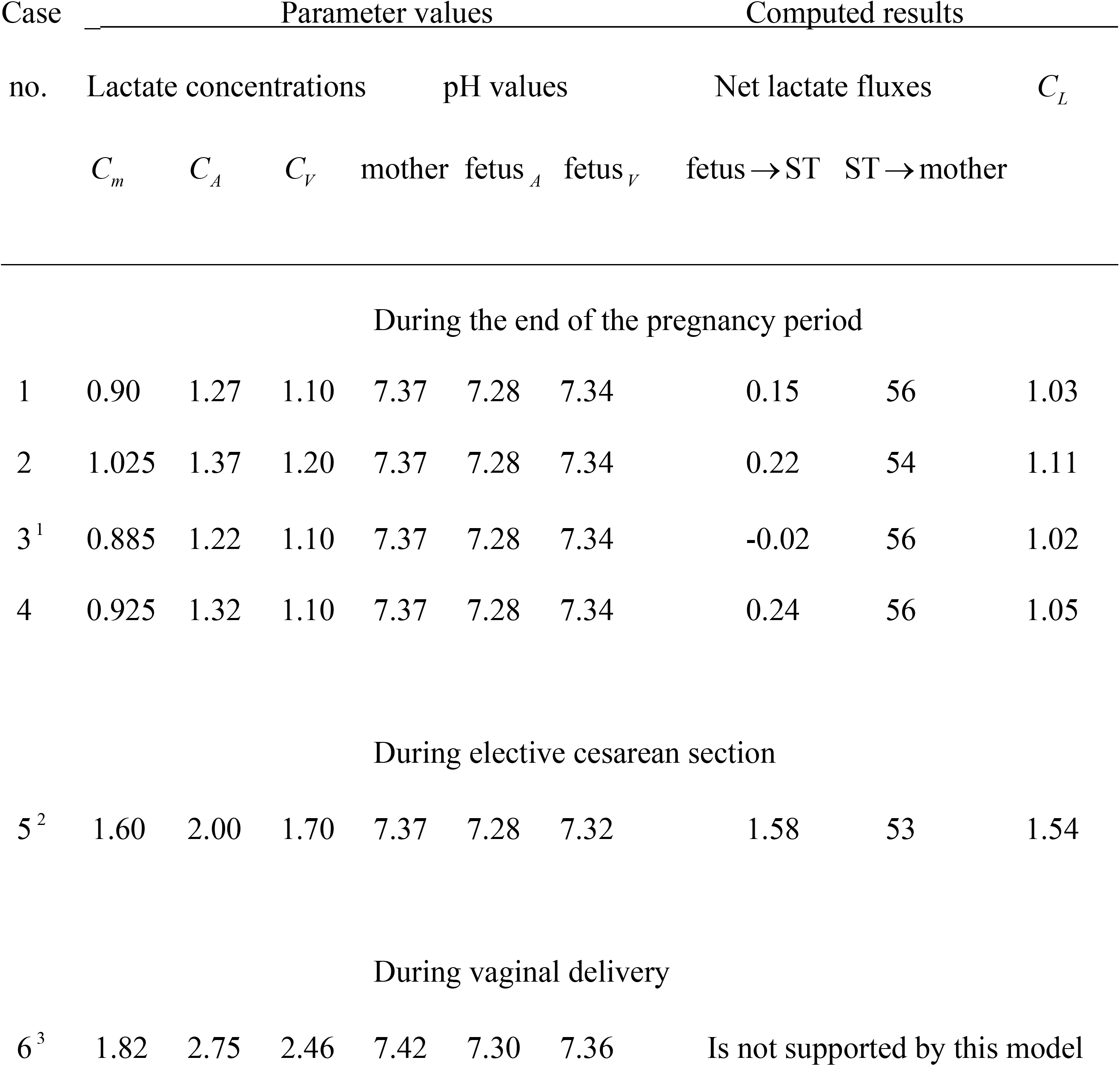
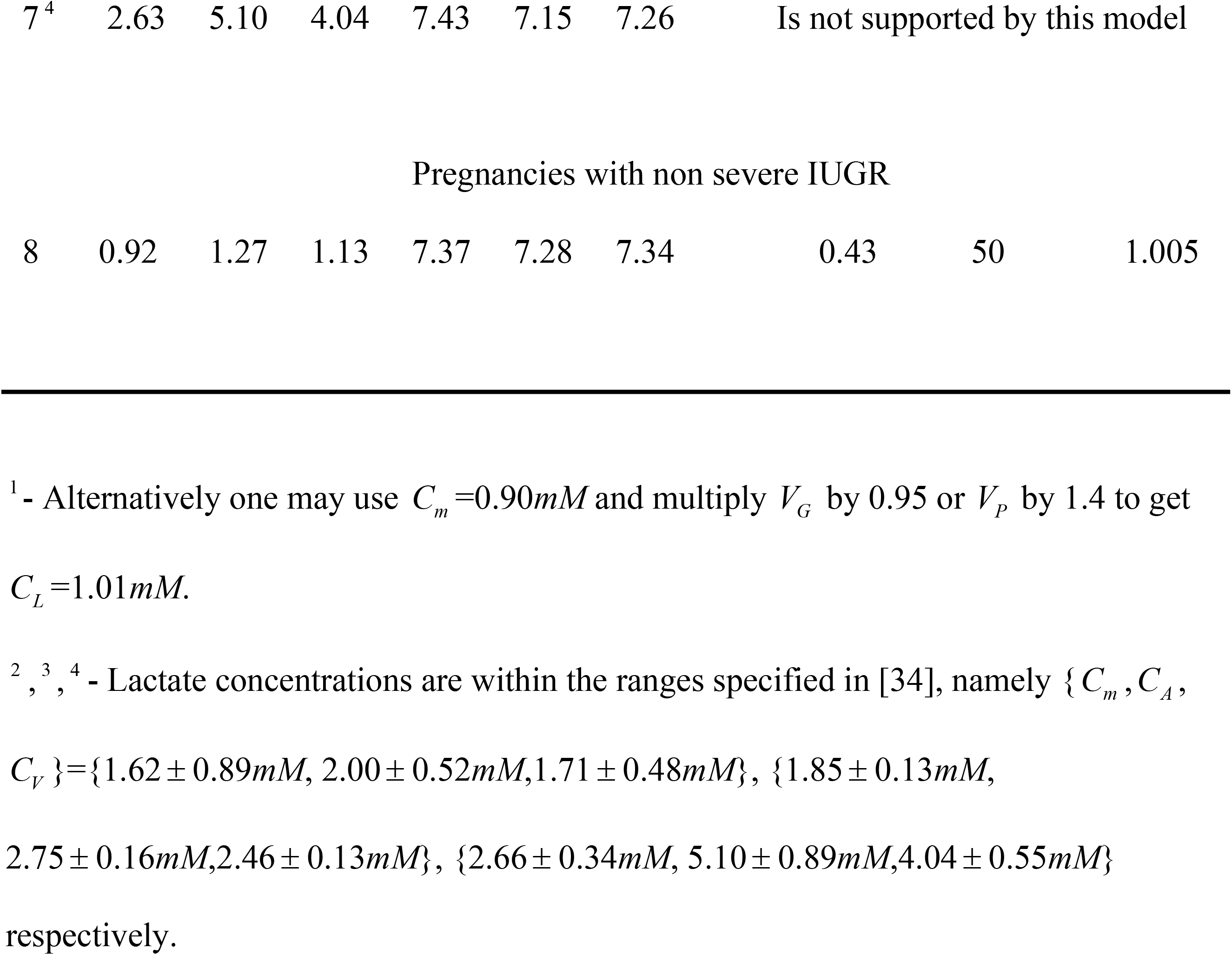
Lactate transport regimes in the uteroplacental unit. Concentrations are given in *mM*, fluxes-in10^−5^*moles* / *s*. Lines 1-7: *P_b_* = 2.4 ×10^6^ *cm*^3^ / *sM*, *P_m_* =12 *P_b_*, *K_p_* = 0.013*M*, *V_G_* =2.5 ×10^−10^ *M* / *s*, other parameter values are specified in Table 1. Parameters for line 8 are specified in S3 Appendix.

When the gradient between lactate levels in fetal arteries and veins changes, the fluxes to and from the placenta are adapted accordingly; a lower fetal lactate production rate is linked to lower maternal lactate level, *C_m_* and/or a lower availability of lactate in the placenta due to a reduced placental glycolysis (a bit lower *V_G_* and/or a considerably higher *V_P_*). All these possible scenarios involve a slower fetal lactate evacuation via its arterioles and slightly lower lactate supply via the venoules to the extent that there is a net lactate supply to the fetus (3^rd^ line, Table 2). Thus, the placenta again serves as a buffer that tries to compensate for lactate shortage in fetal circulation. Opposite effects are demonstrated when fetal lactate production is enhanced (4^th^ line, Table 2).

During labor, lactate production is dramatically increased, leading to higher lactate levels and lower pH values [34, 28]. Line 5 in Table 2 demonstrates how the fetus copes with the elevated lactate levels during an elective cesarean section, enhancing the lactate’s evacuation to the placenta by an order of magnitude. However, during spontaneous labor, contractions significantly reduce blood flow in the uteroplacental unit, glucose concentrations considerably change [35] and metabolic activity is unstable. Indeed, we compute a similar, instead of opposite, sign of flow to and from the fetus in lines 6&7 of Table 2 - an indication that under our model’s assumptions the measured lactate levels cannot be sustained; those are unsteady concentrations inflicted by the turmoil in the uteroplacental unit during the spontaneous labor.

The case of a pregnancy that is complicated by an intrauterine growth restriction, IUGR necessitates utilizing a different set of parameter values due to different morphology, membranes’ structure and characteristics. In S3 Appendix we extend on the implications of this abnormality and specify the relevant parameter values. Our simulation, for the first time, demonstrates the effect of this pathology on lactate fluxes in the uteroplacental unit. As is shown in line 8 of Table 2, although non severe IUGR has a minor effect on lactate concentrations in all the components of the uteroplacental unit [30], it does involve a considerably higher lactate evacuation from the fetus that probably helps to avoid an acidosis.

### B. Determination of physiologically legitimate ranges of parameter values

Many combinations of parameter values, when applied to our simulation, result with unidirectional flows in the fetal arterioles and venoules- an impossible situation that leads to a rejection of those sets, declaring them as non-physiologic ones. For example, multiplying *V_G_* by 1.2 thus enhancing the placental glycolysis rate while keeping all other parameter values intact, leads to such an outcome. This implies that higher *V_G_* values are accompanied by altered maternal and/or fetal lactate concentrations. The interrelation between possible values of several parameters might be demonstrated by replacing the set of 4 parameters used for the computation of Tab. 2 and Fig 3 by the following feasible sets:{*P_b_*, *P_m_*, *K_p_*,*V_G_*} = {2.3×10^6^ *cm*^3^ (*sM*)^−1^,14*P_b_*, 0.015*M*, 2.7 ×10^−10^*Ms*^−1^} or {2.5×10^6^ *cm*^3^ (*sM*)^−1^,10*P_b_*, 0.014*M*, 2.2 ×10^−10^*Ms*^−1^}. Plentiful of other legitimate combinations exist. The above replacements enhance/reduce lactate flow to maternal circulation by about 15% but have no effect on lactate fluxes to and from the fetus. Therefore, measuring lactate flow to the mother will enable a determination of a trustable combination of parameter values. In light of the scarcity of relevant measurements we have no choice but to settle for the ranges of values that appear in Table 1. Any new substantiated parameter value will necessarily narrow the legitimate ranges of other parameter values.

### C. Estimation of the relative importance of the involved mechanisms

Substitution of our solution in the three expressions that constitute the right hand side of Eq.1 enables estimation of the relative contribution of the three mechanisms that govern lactate concentration within the ST. It turns out that the first two terms, the diffusion and the glycolysis, have a similar contribution while lactate degradation is less important but still-significant. The comment to the third line in Table 2 demonstrates how glycolysis within the placenta plays a critical role since diminishing *V_G_* by 5% has a similar effect as raising *V_P_* by 40% in agreement with McNanley & Woods [22] who stated that just a small portion of lactate is consumed within the placenta. A similar (albeit, less pronounced) discrepancy exists in relation to sensitivities with respect to *K*_*G*_ *K_P_*: increasing *K_G_* by 10% is equivalent to reducing *K_P_* by 30%. We conclude that the sensitivity of lactate fluxes with respect to placental glycolysis rate is very high. Another “major player” is the permeability of the MVM. We found that increasing *P_b_* by 10% results in the same increment of the net flow at the BM while placental lactate level, and therefore lactate clearance to the mother, hardly changes. However, increasing *P_m_* by 10% leads to a lower placental lactate concentration and a 350% higher net flow at the BM. Gauging the apparent permeability of the MVM with respect to lactate was quite tricky here as no straightforward measurements are available. Unless the ratio between the weights of the MVM and the BM is measured, using the relative abundance of the transporters on the membranes as a key to determine *P_m_* / *P_b_* might involve significant inaccuracies.

Due to the high variability of normal placentas’ morphology, it is important to have a sensitivity analysis with respect to the geometrical parameter values *S_A_* and *x*_1_. We found that leaving lactate concentrations unchanged while *S_A_* increases, induces enhanced net flux at the BM and higher lactate concentration at the ST. It implies utilization of a significantly lower *V_P_* value or alternatively-both higher *P_m_* and *V_G_* values. Reducing *x*_1_ as well is accompanied by an enhanced net flux at the BM and implies utilization of higher *V_G_* and slightly slower degradation rate. Nevertheless, these scenarios involve parameter values that are within the range specified in Table 1 indicating the reliability of our calibration of parameter values.

## Discussion

In 1993 Schaefer et al [3] suggested a hypothetical lactate cycle in the uteroplacental unit but it ignored the glycolysis in the placenta that was shown to play a major role in ample measurements [1, 28, 30]. No other updated models are known. Moreover, the scarce available measurements relate to concentration values [30, 34] and not to measurements of fluxes.

Here, a mathematical model of the mechanisms that govern lactate transport in the placenta is formulated in order to provide novel insights as well as verifications and explanations for previous observations. The effect of various conditions that prevail in the uteroplacental unit on local lactate fluxes is being fathomed for the first time. This model demonstrates the important, active role that the placenta plays, serving as a lactate buffer by shielding the fetus in case of elevated maternal lactate concentration on the one hand and by enhancing its own metabolic activity in order to supply it to a fetus with low lactate levels on the other hand.

The following assumptions underlie this model:

- 1D simulation is good enough. Previous 3D models for nutrients’ transport via the placenta yielded concentration profiles that were basically 1D. This and the highly complicated and personalized morphology of the human placenta turn 1D models to be a prevalent choice.
- Lactate production due to amino acids e.g. alanine metabolism [2, 36] might be ignored within the placenta.
- Lactate transport across the membranes might be represented by action of the MCT transporters alone, neglecting passive diffusion [11] and activity of other transporters [7, 16], both known to be of ancillary importance.
- Michaelis-Menten kinetics may describe the metabolic reactions. This is the conventional approximation for those complex mechanisms but necessarily it involves inaccuracies.
- The presence of several, other than hydrogen, ions e.g., Na^+^, K^+^ which are known to affect the permeability of the membranes with respect to lactate [13, 16, 29] might be ignored.

As the above simplifications follow the conventions in this field, the reliability of this simulation depends mainly on accurate calibration of the parameter values. In order to validate our calibration we run our simulation for the case of IUGR, using data from two independent laboratories that estimated its effect on lactate concentrations in the uteroplacental unit and on transporters activity, see S3 Appendix. The computations for the effect of the IUGR fully corroborate our approach.

Our results indicate that:

1. Lactate flux to the maternal circulation is about two orders of magnitude higher than the fluxes to and from the fetus, in accord with very early perfusion experiments that succeeded to detect just lactate transport from placenta to mother [1]. Under normal conditions lactate is evacuated from fetal arteries in a rate that outpaces lactate uptake via the umbilical vein by 15-20% approximately. The computed minor lactate supply to the fetus agrees with Suidan et al. [28] findings that the lion share of the fetal lactate originates in self production. A change of the prevailing conditions in any of the involved compartments may convert lactate net clearance to lactate uptake by the fetus. In IUGR complicated pregnancies placental lactate supply to the fetus diminishes.
2. The glycolysis within the placenta is a very important source for lactate and its characteristics must be carefully measured. By contrast, lactate degradation in the ST plays a moderate role thus its rate, presently unknown, might be roughly approximated.
3. Our simulation directs the experimentalists to measurements that are supposed to improve the affectivity of this model, e.g. for want of laboratory data we had to implicitly infer the permeabilities of the confining membranes to lactate. The permeability of the BM that is computed by converging two different methods for computation of the lactate flux there; On the one hand it is the product of the volume of blood that is exchanged between the placenta and the fetus and the umbilical venoarterial lactate concentration difference, Eq.4. On the other hand it is the product of the permeability and the difference between the densities of the occupied binding sites at the two faces of the membrane, Eq.3. However, the permeability of the MVM is the more crucial factor in determining lactate fluxes and in order to get a more precise estimation of it one should know the ratio between the weights of the two confining membranes.
4. During elective cesarean section, parameter values within the uteroplacental unit are being considerably changed. However, as the same mechanisms that prevail during the late pregnancy period are still active, the situation can be simulated by our model. We found an order of magnitude enhancement of activity at the fetal side of the uteroplacental unit with just minor changes at the maternal side. Our inability to simulate vaginal deliveries is an indication of their chaotic features as far as lactate is concerned.

The successful implementation of this simulation for the IUGR pathology indicates at the potential of implementing this model for other pathologies as well, helping to decipher their effect on the lactate transport, once the relevant parameter values are measured. This model does not account for the mutual, dynamic interaction between pH and lactate levels in the plasma. Once measurements yield data regarding the interplay between pH value and lactate level, a more comprehensive model could be formulated.

## Supporting information

**S1 Appendix. Formulation of the boundary conditions**.

**S2 Appendix. Determination of the parameter values**.

**S3 Appendix. Parameter values for the IUGR case**.

## References

1. Hauguel S, Challier JC, Cedard L, Olive G. Metabolism of the human placenta perfused in vitro: glucose transfer and utilization, *O*_2_ consumption, lactate and ammonia production. Pediatr. Res. 1983;17: 729–732.

2. Medina JM, Fernandez E, Bolanos JP, Vicario C, Arizmendi C. Fuel supply to the brain during the early postnatal period. in: Cuezva JM et al, editors., Endocrine and biochemical development of the fetus and neonate. Plenum Press; 1990, p. 175–194.

3. Schaefer A, Piquard F, Dellenbach P, Haberey P. Placenta-fetal “Alanine-lactate cycle” in the human during late gestation. Troph. Res. 1993;7: 103–114.

4. Nagai A, Takebe K, Nio-Kobayashi J, Takahashi-Iwanaga H, Iwanaga T. Cellular expression of the monocarboxylate transporter (MCT) family in the placenta of mice. Placenta 2010;31: 126–133.

5. Moore NP, Picut CA, Charlap JH. Localisation of lactate transporters in rat and rabbit placentae, Int. J. Cell Biol. 2016. Available from: https://doi.org/10.1155/2016/2084252

6. Benirschke K, Burton GJ, Baergen RN. Pathology of the human placenta, 6th ed., Springer; 2012.

7. Inuyama M, Ushigome F, Emoto A, Koyabu N, Satoh S, Tsukimori K, Nakano H, Ohtani H, SawadaY. Characteristics of L-lactic acid transport in basal membrane vesicles of human placental syncytiotrophoblast. Am. J. Cell Physiol. 2002;283: C822–C830.

8. Settle P, Mynett K, Speake P, Champion E, Doughty IM, Sibley CP, D’Souza SW, Glazier J. Polarized lactate transporter activity and expression in the syncytiotrophoblast of the term human placenta. Placenta 2004;25: 496–504.

9. Barta E, Drugan A. A theoretical model of glucose transport suggests symmetric GLUT1 characteristics at placental membranes. J. Membr. Biol. 2014;247(8): 685–694.

10. Plitman MR, Charnock-Jones DS, Burton GJ, Oyen ML. Three-dimensional modeling of human placental terminal villi. Placenta 2016;43: 54–60.

11. Li XB, Gu JD, Zhou QH. Review of aerobic glycolysis and its key enzymes – new targets for lung cancer therapy. Thorac Cancer. 2015;6(1): 17–24.

12. Sun S, Li H, Chen J, Qian Q. Lactic acid: no longer an inert and end-product of glycolysis. Physiology 2017;32: 453–463.

13. Alonso De La Torre SR, Serrano MA, Alvarado F, Medina JM. Carrier-mediated L-lactate transport in brush-border membrane vesicles from rat placenta during late gestation. Biochem. J. 1991;278: 535–541.

14. Turner RJ. Kineticanalysis of a family of cotransporter models. Biochim. Biophys. Acta 1981;649: 269–280.

15. Sanders D, Hansen U-P, Gradmann D, Slayman CL. Generalized kinetic analysis of ion-driven cotransport systems: a unified interpretation of selective ionic effects on Michaelis parameters. J. Memb. Boil. 1984;77: 123–152.

16. Balkovetz DF, Leibach FH, Mahesh VB, Ganapathy V. A proton gradient is the driving force for uphill transport of lactate in human placental brush-border membrane vesicles. J. Biol. Chem. 1988;263(27): 13823–13830.

17. Barta E. Transport of docosahexaenoic acid via the human placenta: a theoretical study. J. Membr. Biol. 2019;252: 617–626.

18. Pfeuffer J, Tkác I, Gruetter R. Extracellular-intracellular distribution of glucose and lactate in the rat brain assessed noninvasively by diffusion-weighted IH nuclear magnetic resonance spectroscopy in vivo. J Cereb Blood Flow Metab. 2000;20(4): 736–746.

19. Al-Husari M, Webb SD. regulation of tumor intracellular pH: A mathematical model examining the interplay between H^+^ and lactate. J. Theor. Biol. 2013;322; 58–71.

20. Bittner CX, Loaiza A, Ruminot I, Larenas V, Sotelo-Hitschfeld T, Gutiérrez R, Córdova A, Valdebenito R, Frommer WB, Barros LF. High resolution measurement of the glycolytic rate. Front. Neuroenergetics 2010. Available from: https://doi:10.3389/fnene.2010.00026

21. Michelsen TM, Holme AM, Holm MB, Roland MC, Haugen G, Powell TL, Jansson T, Henriksen T. Uteroplacental glucose uptake and fetal glucose consumption: a quantitative study in human pregnancies. J. Clin. Endocrinol. Metab. 2019;104: 873–882.

22. McNanley T, Woods J. Placental physiology, Glob. libr. women’s med., (ISSN: 1756-2228) 2008. Available from: https://doi:10.3843/glowm.10195

23. Talaiezadeh A, Shahriari A, Tabandeh MR, Fathizadeh P, Mansouri S. Kinetic characterization of lactate dehydrogenase in normal and malignant human breast tissues. Cancer cell International 2015. Available from: https://doi:10.1186/s12935-015-0171-7

24. Lambeth MJ, Kushmerick MJ. A computational model for glycogenolysis in skeletal muscle. Ann. Biomed. Eng. 2002;30: 808–827.

25. Cowley EA, Sellers MC, Illsley NP. Intracellular pH homeostasis in cultured human placental syncytiotrophoblast cells: recovery from acidification. Am. J. Physiol. Cell Physiol. 2005;288(4): C891–C898.

26. Soma-Pillay P, Nelson-Piercy C, Tolppanen H, Mebazaa A. Physiological changes in pregnancy. Cardiovasc J. Afr. 2016;27(2): 89–94.

27. Cao Y, Wang M, Yuan Y, Li C, Bai Q, Li M. Arterial blood gas and acid-base balance in patients with pregnancy-induced hypertension syndrome. Experimental and therapeutic medicine 2018. Available from: https://doi.org/10.3892/etm.2018.6893

28. Suidan JS, Antoine C, Silverman F, Lustig ID, Wasserman JF, Young BK. Human maternal-fetal relationships. J. Perinat. Med. 1984;12: 211–217.

29. Araos J, Silva, Salsoso R, Sa’ez T, Barros E, Toledo F, Gutie’rrez J, Pardo F, Leiva A, Sanhueza C, Sobrevia L. Intracellular and extracellular pH dynamics in the human placenta from diabetes mellitus. Placenta 2016;43: 47–53.

30. Marconi AM, Paolini CL, Zerbe G, Battaglia FC. Lactacidemia in intrauterine growth restricted (IUGR) pregnancies: relationship to clinical severity, oxygenation and placental weight. Pediatr. Res. 2006;59(4): 570–574.

31. Mielke G, Benda N. Cardiac output and central distribution of blood flow in the human fetus. Circulation 2001;103: 1662–1668.

32. Fineman JR, Clyman R. Fetal Cardiovascular Physiology. In: Creasy & Resnik’s Maternal Fetal Medicine. 7th ed., Elsevier Publ.: 2014, pp. 146–155

33. Jimenez V, Henriquez M, Llanos P, Riquelme G. Isolation and purification of human placental plasma membranes from normal and pre-eclamptic pregnancies. A comparative study. Placenta 2004;25: 422–437.

34. Schneider H, Danko J, Huch R, Huch A. Homeostasis of fetal lactate metabolism in late pregnancy and the changes during labor and delivery. Europ. J. Obstet. Gynec. Reprod. Biol. 1984;17: 183–192.

35. Maheux PC, Bonin B, Dizazo, Guimond P, Monier D, Bourque J, Chiasson J. Glucose homeostasis during spontaneous labor in normal human pregnancy. J. Clin, Endocrinol. Metab. 1996;81: 209–215.

36. Holzman IR, Philipps AF, Battaglia FC. Glucose metabolism, lactate, and ammonia production by the human placenta in vitro. Pediat. Res. 1979;13: 117–120.

37. Settle P, Sibley CP, Doughty IM, Johnston T, Glazier JD, Powell TL, Jansson T, D’Souza SW. Placental lactate transporter activity and expression in intrauterine growth restriction. J. Soc. Gynecol. Investing. 2006;13: 357–363.

38. Sibley CP, Turner MA, Cetin I, Ayuk P, Boyd CAR, D’Souza SW, Glazier JD, Greenwood SL, Jansson T, Powell TL. Placental Phenotypes of Intrauterine Growth. Pediatr. Res. 2005;58: 827–832.

